# Oxidation of ΔFOSB at Cys172 Controls Hippocampal Gene Targets and Learning

**DOI:** 10.1101/2025.10.14.682315

**Authors:** Haley M. Lynch, Daniela Anderson, Brandon Hughes, Galina Aglyamova, Sam Yeh, Yoshinori N. Onishi, Molly Estill, Bryan Granger, Hannah M. Cates, Stefano Berto, Jeannie Chin, Li Shen, Eric J. Nestler, Gabby Rudenko, A.J. Robison

**Author notes:** These authors contributed equally to this work (Haley M. Lynch, Daniela Anderson).

## Abstract

Imbalance of reduction/oxidation (redox) in the brain is associated with numerous diseases including Alzheimer’s disease (AD), substance abuse disorders, and stroke. Moreover, cognitive decline can be caused by neuronal dysfunction that precedes cell death, and this dysfunction is in part produced by altered gene expression. However, the mechanisms by which redox state controls gene expression in neurons are not well understood. ΔFOSB is a neuronally enriched transcription factor critical for orchestrating gene expression underlying memory, mood, and motivated behaviors. It is dysregulated in many conditions including AD. We showed recently that ΔFOSB forms a redox-sensitive disulfide bond between cysteine 172 (C172) of ΔFOSB and C279 of its preferred binding partner JUND. This bond works as a redox switch to control DNA-binding, based on studies of recombinant proteins *in vitro*. Here, we show that this redox control of ΔFOSB function *in vitro* is conserved *in vivo*. We show that ΔFOSB C172 forms a redox-sensitive disulfide bond with JUND that regulates the stability of this AP1-transcription factor complex and its binding to DNA in cells. We also validate the formation of ΔFOSB-containing complexes held together via disulfide bonds in mouse brain *in vivo*. We show that exogenous oxidative stress reduces ΔFOSB binding to gene targets in mouse brain and that *Fosb* C172S knock-in mice, which lack a functional ΔFOSB redox switch, are insensitive to this oxidation-dependent reduction in target gene binding, demonstrating that ΔFOSB is regulated by a redox switch that modulates binding to target genes in the hippocampus. Finally, we demonstrate that *FosB* C172S knock-in mice are less sensitive to cognitive dysfunction induced by oxidative stress. This evidence supports ΔFOSB as an important mediator of oxidative stress-driven changes in gene expression and cognition and implicates ΔFOSB as a possible therapeutic target for diseases associated with oxidative stress in the brain, including AD.

## Introduction

From psychiatric to neurodegenerative disorders, oxidative stress is a common driver of neurological and psychiatric dysfunction [1]. Oxidative stress is characterized by an imbalance of reactive oxygen species (ROS) and the antioxidants and metabolic processes that work to neutralize them. This excess of ROS can come from a variety of sources including external stimuli like drugs of abuse or toxins [2], and internal causes such as dysfunctional mitochondria and decreases in antioxidant levels that accompany the aging process [3]. Oxidative stress leads to an array of problems such as lipid peroxidation, dysfunctional DNA repair and replication, malfunctioning mitochondria, and inflammation, all of which accompany disease pathogenesis including that seen in Alzheimer’s disease (AD) [4]. Another characteristic of such reduction-oxidation (redox)-sensitive conditions is alteration in gene expression [5]; however, whether and how oxidative stress drives these changes in gene expression remains elusive.

ΔFOSB is a neuronally enriched transcription factor critical for orchestrating gene expression underlying memory, mood, and motivated behaviors [6, 7]. ΔFOSB is induced by a variety of stimuli including chronic exposure to drugs of abuse [8, 9], seizures [10], stress [11, 12], and learning [13], among other perturbations. Additionally, it is upregulated in the hippocampus of rodent models of AD and post-mortem hippocampi collected from patients with AD and may serve to protect against seizures in AD models [14, 15]. ΔFOSB is a product of the *Fosb* gene that is prematurely truncated due to splice variation, and the resulting removal of degron domains in the C terminus provides it with unique stability compared to full-length FOSB and all other FOS family proteins. This stability allows it to accumulate with chronic stimulation and remain elevated in the brain for weeks after the final stimulus [8, 16-18]. As a result, ΔFOSB is thought to mediate long-term changes in gene expression as seen in chronic diseases like addiction or AD [6, 7].

ΔFOSB regulates gene expression by binding to DNA at specific AP-1 consensus sequences (TGA C/G TCA). As an alternative splice variant of *Fosb*, ΔFOSB contains a disordered N-terminal domain (residues Met^1^-Glu^156^) and a bZIP domain composed of a basic region (Lys^157^-Arg^177^) that contains a DNA-binding motif and a leucine zipper (Thr^180^-His^218^) which forms a coiled-coil with a dimerization partner [18]. ΔFOSB dimerizes predominantly to JUND, another bZIP protein, to form an activator-protein-1 (AP-1) transcription factor. Complex formation occurs upon dimerization of the leucine zippers of ΔFOSB and JUND, clamping the basic regions on either side of the DNA strand into the major groove like a pair of forceps (**Fig. 1A**). This complex binds AP-1 sites that are found at both promoter regions [19] as well as putative enhancer regions in gene bodies and intergenic regions [20] associated with target genes. Although JUND appears to be the primary binding partner *in vivo* [21, 22], *in vitro* evidence suggests ΔFOSB homodimers may also exist and bind to DNA [23, 24]. Once bound to DNA, ΔFOSB can either repress or activate its gene targets [25].

**Figure 1.**
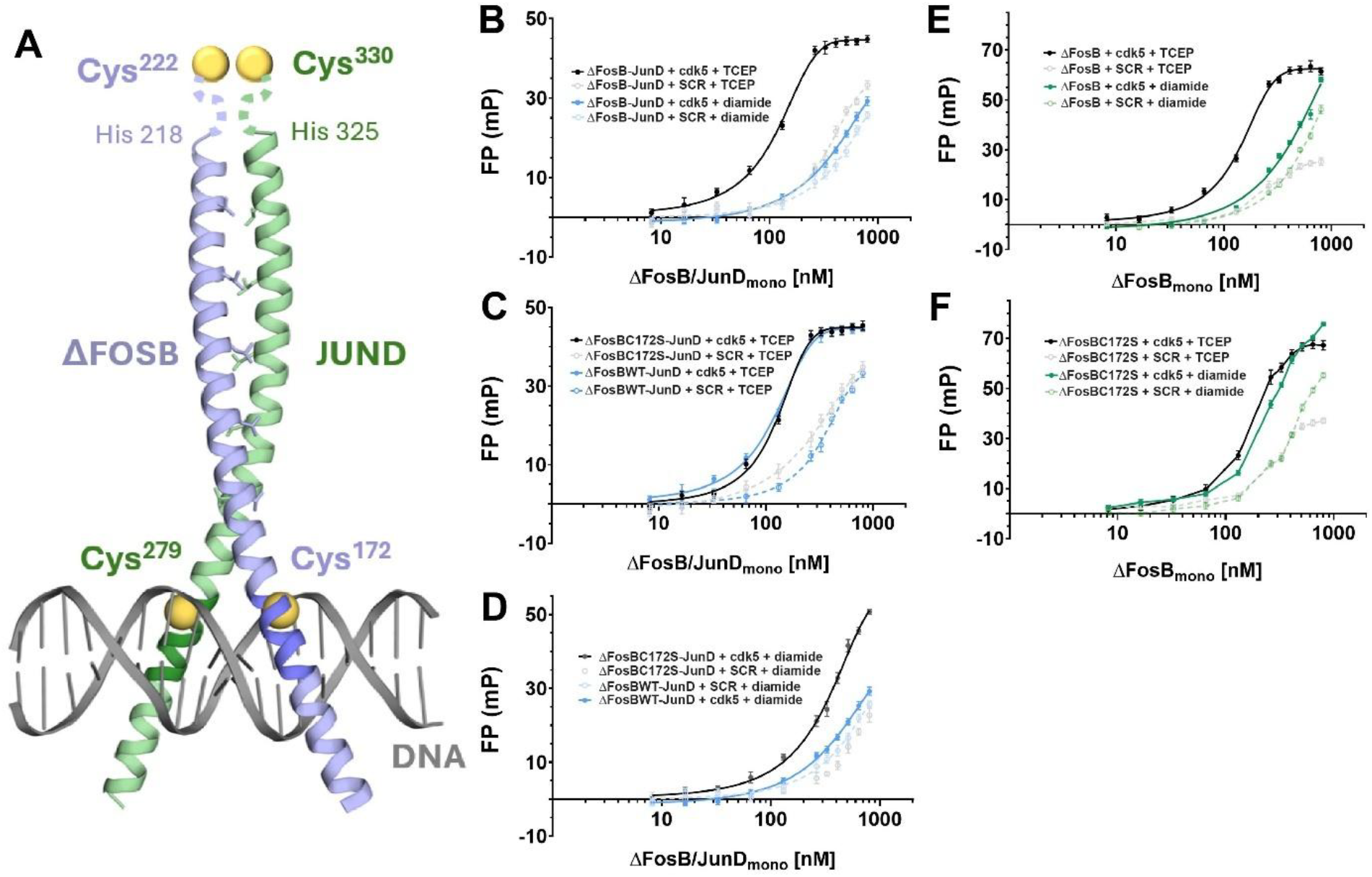
ΔFOSB C172 mediates oxidation-dependent inhibition of DNA binding *in vitro*. **A)** Structure of ΔFOSB/JUND complex bound to DNA illustrating the juxtaposition of ΔFOSB C172 and JUND C279 and proximity to the DNA binding regions (mouse amino acid numbers). **B)** FP assay using purified ΔFOSB/JUND complexes shows oxidizing reagents reduce binding to the *Cdk5* AP-1 site to levels similar to scrambled DNA. **C)** C172S mutation has no effect on specific DNA binding under reducing conditions. **D)** C172S mutation prevents the oxidation-dependent reduction in DNA binding seen in WT ΔFOSB. **E)** ΔFOSB homodimer binding to the *Cdk5* AP-1 site is redox-sensitive. **F)** C172S mutation prevents redox sensitivity of ΔFOSB homodimer binding to DNA. *NB:* Panels B, C, and D show the same data for wild-type ΔFOSB/JUND binding for comparison to mutant under different conditions.

Recent i*n vitro* data from our laboratories suggest that ΔFOSB is sensitive to redox state. That is, ΔFOSB contains redox-sensitive intermolecular disulfide bonds that occur via oxidation of cysteine residues found in ΔFOSB and its partner protein [26]. ΔFOSB has a total of four cysteine residues; two in the disordered region C15 and C44 (both unstudied), one near the leucine zipper region C222 [27], and the fourth and most studied in the DNA binding motif, C172 [27, 28]. C172 behaves as a “redox-switch” to control DNA binding under different redox conditions *in vitro* [28], whereas C222 may act in a “zip-lock” fashion to stabilize the leucine zipper interaction in a ΔFOSB complex [27]. This evidence of redox-sensitivity was found using recombinant protein purified from insect cells, and as of yet, there is no evidence as to whether or not this effect is conserved *in vivo*. Moreover, evidence is lacking as to whether ΔFOSB could be driving pathological changes in gene expression as a consequence of the brain being in a state of oxidative stress, as in AD.

To test the possibility of ΔFOSB’s role as a mediator of oxidative stress and downstream alterations in gene expression in diseases like AD, we sought to mimic a state of oxidative stress through chemical treatment of Neuro2a (N2a) cells, a neuroblastoma cell line, or hippocampal mouse brain tissue with redox state modifying agents to see the effects of oxidative stress on ΔFOSB’s ability to bind JUND and to bind DNA. Additionally, we manipulated ΔFosB at cysteine residues C172 and/or C222 to determine their redox sensitivity and their impact on how ΔFOSB responds to oxidative stress. Our results reveal that oxidative stress promotes the formation of ΔFOSB complexes, however, it also reduces its ability to bind DNA *in vivo*. This attenuation of ΔFOSB DNA binding renders the complex transcriptionally inactive, which agrees with previous *in vitro* studies. We further used two novel transgenic mouse lines (*Fosb* C172S knockin mouse and HA-tagged *Fosb* mouse) to establish that similar redox-dependent complexes can form in the mouse brain and that ΔFOSB C172 oxidation regulates DNA binding in the hippocampus. Finally, we show that ΔFOSB C172 oxidation inhibits some forms of hippocampus-dependent learning, indicating that oxidation of ΔFOSB at this site is a critical mechanism that regulates cognition and could potentially connect oxidative stress to cognitive decline in the early stages of AD.

## Methods

See Supplemental Materials

## Results

### Cysteine 172 oxidation regulates ΔFOSB binding to DNA *in vitro*

Because ΔFOSB C172 forms a disulfide bridge with JUND C279 which reduces binding to DNA in biochemical assays [28, 29] (**Fig 1A**), we set about determining the role of the ΔFOSB/JUND redox switch in more physiologically relevant settings. First, to determine whether oxidative conditions actually prevent ΔFOSB/JUND complexes from binding to the AP-1 consensus sequence from a known target gene, we incubated purified full-length ΔFOSB/JUND heterodimers with a double-stranded oligonucleotide containing the AP-1 consensus sequence in the *Cdk5* gene promoter and measured DNA binding using a fluorescence polarization (FP) binding assay under oxidizing and reducing conditions. We found that in the presence of Tris(2-carboxyethyl)phosphine (TCEP), a potent reducing agent, the ΔFOSB/JUND complex bound the *Cdk5* oligonucleotide with strong specificity compared to the scrambled control oligonucleotide (**Fig 1B**). However, when we used diamide to cause oxidation and disulfide bond formation, we found that the ΔFOSB-JUND complex had greatly reduced binding to *Cdk5* DNA and was also no longer able to distinguish between the *Cdk5* and scrambled oligonucleotides (**Fig 1B**).

To assess whether this redox sensitivity of the ΔFOSB/JUD complex DNA binding is driven by C172, we mutated cysteine 172 to serine (C172S), which conserves the general size of the side chain, but does not allow oxidation and disulfide bond formation. Using the FP assay, we found that under reducing conditions the ΔFOSBC172S/JUND complex bound to *Cdk5* DNA with similar affinity to that of the wild type (WT) ΔFOSB/JUND complex and discriminated between the *Cdk5* and scrambled oligonucleotides similarly, indicating that both the WT and mutant AP1 transcription factors bound DNA with comparable specificity (**Fig 1C**). However, under oxidizing conditions (i.e., addition of diamide), the ability to specifically bind to the *Cdk5* oligonucleotide was, to a large extent, preserved for ΔFOSB C172S/JUND heterodimers but lost for WT ΔFOSB-JUND (**Fig 1D**). Thus, oxidation of ΔFOSB C172 is sufficient to trigger the disruption of DNA binding both in terms of DNA binding affinity and specificity.

Our previous work has also shown that purified ΔFOSB can form homodimers *in vitro* [23, 30]. To determine whether such prospective homomers can bind to DNA as well and whether that DNA binding is redox-sensitive, we performed the same FP binding assay using purified full-length ΔFOSB homomers. We found that ΔFOSB homodimers bind to the *Cdk5* oligonucleotide with much greater affinity than to the scrambled control oligonucleotide under reducing conditions (just like ΔFOSB/JUND heterodimers), whereas under oxidative conditions ΔFOSB binds DNA overall with less affinity and it binds to both the *Cdk5* and scrambled oligonucleotides similarly (**Fig 1E**). Consistent with the critical role of ΔFOSB C172 in placing DNA-binding of heterodimers under redox control, mutant ΔFOSB C172S homodimers bound to the *Cdk5* oligonucleotide similarly under reducing and oxidizing conditions (**Fig 1F**). These data confirm that ΔFOSB C172 acts as a redox switch in both JUND heterodimers and in homodimers to control binding to a target AP-1 sequence *in vitro*.

### Oxidation regulates ΔFOSB oligomerization in cells

To investigate whether ΔFOSB redox sensitivity is preserved in cells, we used the neuron-like Neuro2a mouse neuroblastoma line transfected with HA-ΔFOSB and FLAG-JUND, as we have done previously [27]. In order to determine whether the endogenous redox state of the cell could affect disulfide bond formation in ΔFOSB complexes, we transfected Neuro2a cells with JUND and WT ΔFOSB, then treated the intact cells with increasing concentrations of diamide, an oxidizing reagent (**Fig 2A**). We then probed the cell extracts with Western blot using a FOSB antibody after SDS-PAGE using reducing (with dithiothreitol [DTT] and β-mercapto ethanol) or non-reducing gel conditions. Under reducing SDS-PAGE, we saw the expected ΔFOSB doublet at M_r_ 35/37 kDa, which represent distinct isoforms (presumably different phosophorylation states) of a ΔFOSB monomer. We found that under non-reducing gel conditions, there was accumulation of a ΔFOSB-containing complex around 75-85 kDa and a loss of bands around the typical monomeric M_r_ of 35-37 kDa (**Fig 2B&C**), indicating formation of homomeric ΔFOSB dimers or ΔFOSB-JUND dimers covalently bonded through disulfide bridges in an oxidation-dependent manner in cells (ANOVA F=3.503, p=0.023).

**Figure 2.**
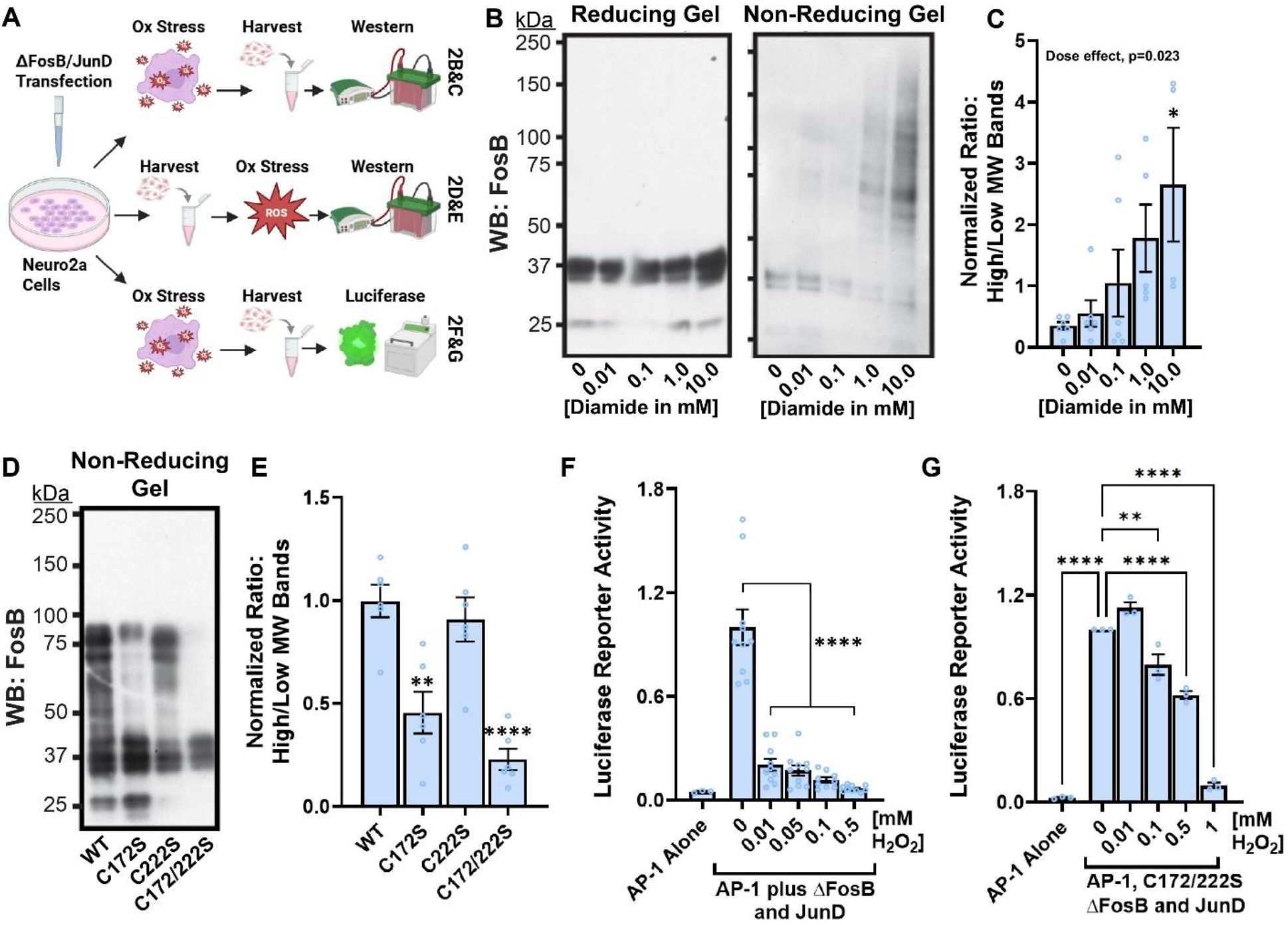
ΔFOSB C172 mediates oxidation-dependent complex formation and AP-1 transactivation in Neuro2a cells. **A)** Schematic of Neuro2a cell transfection, treatments, and measures (Biorender). **B)** Neuro2A cells were transfected with WT ΔFOSB and JUND and exposed to exogenous diamide. Western blot reveals a dose-dependent increase in high molecular weight complexes containing ΔFOSB on non-reducing gels (right) that are absent when reducing reagents are added (left). **C)** Quantification demonstrating dose dependence of oxidative stress-induced ΔFOSB complexes. **D)** Transfected Neuro2a cell homogenates treated with NEM reveal that C172 mutation to serine prevents formation of high molecular weight complexes, quantified in **(E). F)** Luciferase assay in Neuro2a cells transfected with WT ΔFOSB/JUND shows reduced transactivation of AP-1 reporter when cells are treated with oxidizing H_2_O_2_. **G)** Low doses of H_2_O_2_ have little or no effect on C172S ΔFOSB/JUND transactivation of the AP-1 reporter. All graphs depict mean +/-SEM; *:p<0.05, **:p<0.01, ****:p<0.0001.

To determine the role of C172 and C222 in the production of these covalently-linked ΔFOSB/ΔFOSB or ΔFOSB/JUND complexes, we transfected Neuro2a cells with FLAG-JunD and either HA-tagged WT ΔFosB or one of the mutants ΔFosB C172S, ΔFosB C222S, or ΔFosB C172/222S, respectively. Cells were lysed in the presence of N-ethylmaleimide (NEM), an agent that modifies free cysteines to prevent oxidation but allows preservation of existing disulfide bridges [27] (**Fig 2A**). We then performed non-reducing SDS-PAGE and Western blot using an anti-FOSB antibody and observed that WT ΔFOSB produced the expected doublet at molecular weight 35/37 kDa, but also a clear band grouping around 75-85 kDa (**Fig 2D**), consistent with disulfide bonded complexes. ΔFOSB C172S showed less of these high-molecular weight bands (**Fig 2D**, quantified in **2E**), indicating that C172 is critical to form an intermolecular disulfide bond in Neuro2a cells (ANOVA F=17.56, p<0.0001). We further found that the C222S mutation had little effect on high molecular weight complex formation, either in the context of WT C172 or C172S (**Fig 2D&E**).

### Effect of oxidation on C172 ΔFOSB-driven reporter gene expression in cells

As we have shown that ΔFOSB binding to DNA *in vitro* is inhibited by C172 oxidation (**Fig 1**) and that oxidation at this residue occurs in cells (**Fig 2B-E**), we next sought to determine whether ΔFOSB oxidation in cells inhibits the ability of ΔFOSB to drive an AP-1 consensus sequence-containing reporter construct. We transfected Neuro2a cells with HA-ΔFOSB and FLAG-JUND along with a 5xAP-1 luciferase reporter construct, then treated the cells with increasing concentrations of hydrogen peroxide (H_2_O_2_) to drive oxidative stress (**Fig 2A**). We found that oxidative stress strongly reduced ΔFOSB/JUND-driven luciferase reporter activity, even at low concentrations (**Fig 2F**; ANOVA F=56.03, p<0.0001). In order to determine whether cysteine oxidation in ΔFOSB was responsible for this decrease in reporter activity (rather than general effects of oxidation on the reporter, luciferase, or other essential cell machinery), we performed an identical experiment using ΔFOSB C172/222S. Using this construct, we saw little effect of H_2_O_2_ at concentrations that completely abrogated reporter activity with WT ΔFOSB and did not see a complete reduction of reporter activity until H_2_O_2_ reached 1 mM (**Fig 2G**; ANOVA F=236.7, p<0.0001). These data indicate that oxidation of cysteines in ΔFOSB reduces its ability to transactivate AP-1 gene expression in cells.

### ΔFosB oxidation occurs in mouse brain

We next sought to determine whether redox-dependent ΔFOSB complexes occur in mouse brain. Unfortunately, existing antibodies for ΔFOSB produce a non-specific band at 75 kDA (see [31] and **Fig 3A**), exactly the region in which ΔFOSB homodimers and ΔFOSB-JUND heterodimers are found (75-80kDa). Therefore, we created a transgenic mouse with an HA-tag on the N-terminus of the endogenous *FosB* gene (Fosb^HA^) such that all *Fosb* gene products, including full-length FOSB and ΔFOSB, are tagged with HA (**Fig S1**), allowing us to purify ΔFosB-containing complexes from brain and determine their redox-dependent oligomeric state. To ensure that *Fosb* gene products were HA-tagged in the brain, we treated wild type and FosB^HA^ mice with seven days of 20 mg/kg i.p. cocaine to induce *Fosb* gene expression in limibic brain regions, then performed immunofluorescence on brain slices for both ΔFOSB/FOSB and HA (**Fig 3C**). We saw robust expression of ΔFOSB/FOSB in the hippocampus and nucleus accumbens, brain regions known to show cocaine induction of ΔFOSB, and we saw HA expression only in the Fosb^HA^ mice, with a nuclear expression pattern that very strongly overlapped with ΔFOSB/FOSB.

**Figure 3.**
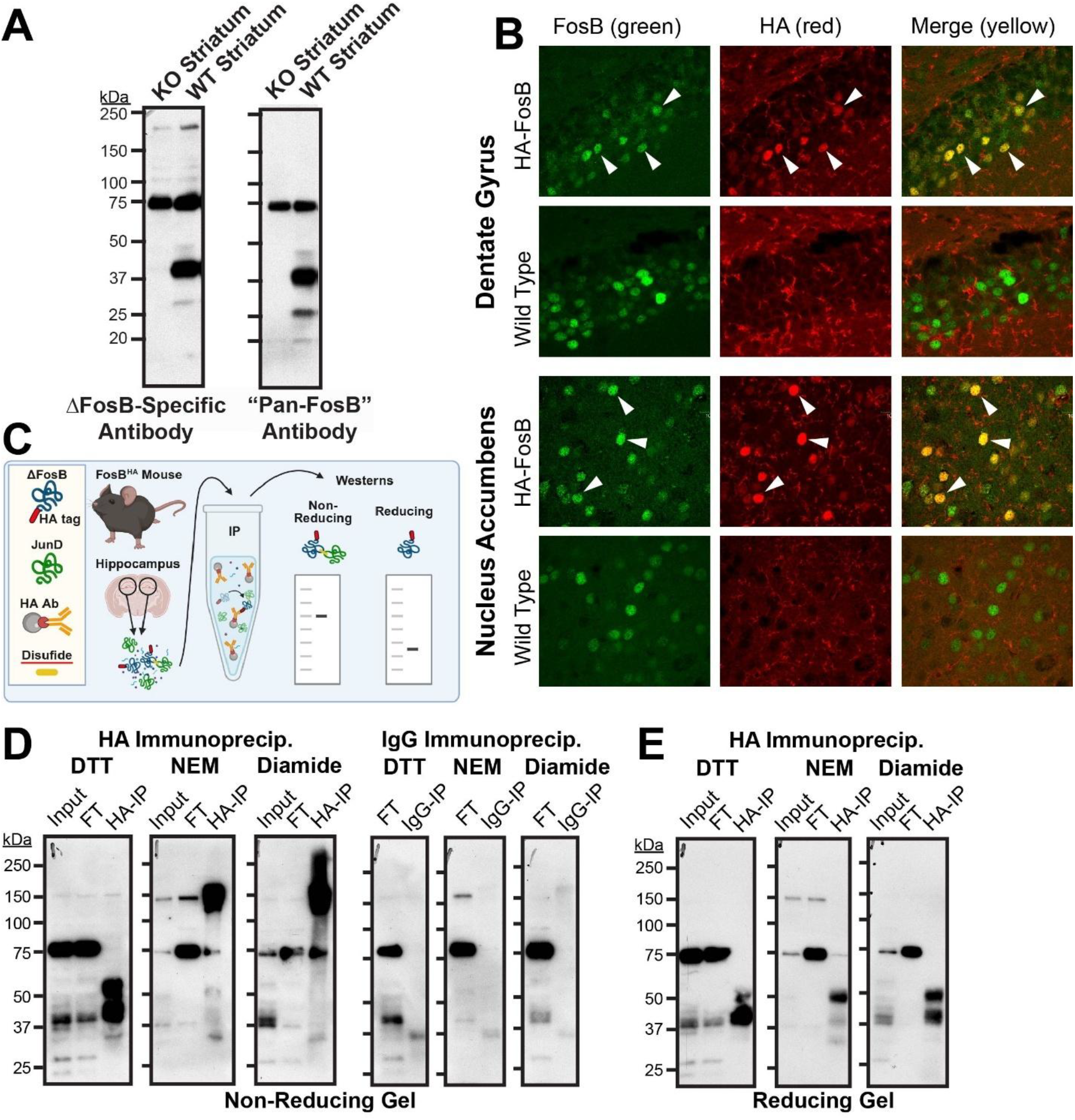
ΔFosB forms redox-dependent complexes in the mouse brain. **A)** Western blot of WT and *Fosb* knockout mouse tissue reveals that existing antibodies consistently detect a non-specific band at 75 kDa that is not a *Fosb* gene product. **B)** Immunofluorescent staining of hippocampus and nucleus accumbens reveals HA signal overlapping FOSB signal in FOSB^HA^ mouse brain, but the absence of an HA signal in WT mouse brain. **C)** Schematic of FOSB^HA^ immunoprecipitation to capture redox-sensitive complexes in mouse hippocampus (Biorender). **D)** Immunprecipitation using an HA antibody in DTT buffer reveals enrichment of monomeric ΔFOS/FOSB at 35/37 and 50 kDa, and the absence of the non-specific 75 kDa band in the IP (but not input or flowthrough – FT). Treatment with NEM or diamide reveal high molecular weight complexes containing ΔFOS/FOSB in the HA-IP at 75 and 150 kDa. All ΔFOSB/FOSB bands are absent in control IgG-IP. **E)** All high molecular weight complexes are absent when reducing agents are added to SDS-PAGE gels.

To determine whether disulfide bonds occur in ΔFOSB complexes in the brain, we again treated FOSB^HA^ mice with seven days of 20 mg/kg i.p. cocaine to induce *Fosb* gene expression, then homogenized hippocampal brain tissue under reducing conditions (with DTT), redox preserving conditions (with NEM), or oxidizing conditions (with diamide). We then performed an immunoprecipitation using the HA antibody or IgG as a control (**Fig 3C**). When we ran the resulting inputs, flowthroughs (FT), or immunoprecipitates (HA-IP) on non-reducing SDS-PAGE and performed Western blot for ΔFOSB/FOSB, we detected enriched ΔFOSB (35/37 kDa) and FOSB (50 kDa) in the reduced DTT HA-IP samples (**Fig 3D**), and these bands were absent in the IgG control IP. In samples homogenized with NEM to preserve redox state, we saw that the ΔFOSB/FOSB immunoreactivity was greatly reduced at 35-50 kDa, but complexes could be seen around 75 and 150 kDa, indicative of ΔFOSB homo- and heterodimers as well as homotetramers, which we have reported before in cells [27]. When oxidation was artificially increased using diamide in the homogenate, we also saw a strong enrichment of the 75 and 150 kDa bands, along with higher molecular weight complexes that could include less physiologically relevant disulfide bonds with other proteins. Critically, all of these higher molecular weight complexes were absent and only the typical monomeric 35-50 kDa ΔFOSB/FOSB bands were present when reducing gels were used (**Fig 3E**). Together, these data indicate that ΔFOSB/FOSB can form redox-dependent higher molecular weight complexes in the mouse brain consistent with disulfide bond formation.

### C172 oxidation regulates ΔFOSB binding to target genes in the brain

Knowing that ΔFOSB can be oxidized in the mouse brain, we next explored whether oxidation at C172 in the brain regulates ΔFOSB binding to gene targets, as it did *in vitro* and in cells. To do this, we generated a transgenic mouse line (FOSB^C172S^) with a point mutation in the *Fosb* gene causing cysteine 172 to be converted to a serine (**Fig S2**), similar to the plasmid constructs used in our cultured cell experiments. These mice had normal body weight and appearance, indicating no gross developmental abnormalities. Moreover, these mice had no differences in elevated plus maze, sucrose preference, novel object discrimination, or cocaine conditioned place preference (**Fig S3**), indicating that this mutation did not affect ΔFOSB-dependent reward or learning behaviors in the absence of exogenous oxidative stress.

We induced systemic oxidative stress in FOSB^C172S^ and WT littermate control mice by treating them with 25 mg/kg/day potassium dichromate (K_2_Cr_2_O_7_; abbreviated KCr) in their drinking water for 7 days, with mice consuming normal drinking water serving as a control. KCr increases protein oxidation in the brain [32]. This treatment did not alter expression of ΔFOS/FOSB protein in the brain (**Fig S4A**), but it did increase oxidation of brain proteins (**Fig S4B**) in WT mice, as expected.

To assess the effects of oxidative conditions in the brain on ΔFOSB binding to gene targets, we harvested dorsal hippocampus from FOSB^C172S^ and WT mice after KCr or control treatment and performed cleavage under targets and release using nuclease (CUT&RUN) on this tissue using a ΔFOSB-specific antibody, as we have previously published [20]. We found that, in WT mice, KCr treatment reduced ΔFOSB-bound peaks by 59% compared to normal water drinking controls (2013 peaks reduced to 820 peaks; **Fig 4A**), indicating that oxidation reduces ΔFOSB DNA binding in the brain. Critically, we found that KCr treatment increased ΔFOSB binding peaks in FOSB^C172S^ mice by 68% (2447 peaks increased to 4101 peaks; **Fig 4A**), suggesting that C172 oxidation is essential for the effects of oxidative stress on ΔFOSB binding to target genes. There was no change in peak width (**Fig 4B**) or general distribution of ΔFOSB sites across genomic features (**Fig 4C**) by genotype or treatment. We likewise found that KCr treatment reduced the number of genes associated with ΔFOSB binding in WT mice by 56% (1544 bound genes reduced to 681), and that this effect was reversed in FosB^C172S^ mice (61% increase, 1824 bound genes up to 2947; **Fig 4D**), indicating that oxidative stress reduces the number of gene targets bound by WT ΔFOSB through oxidation of the C172 residue.

**Figure 4.**
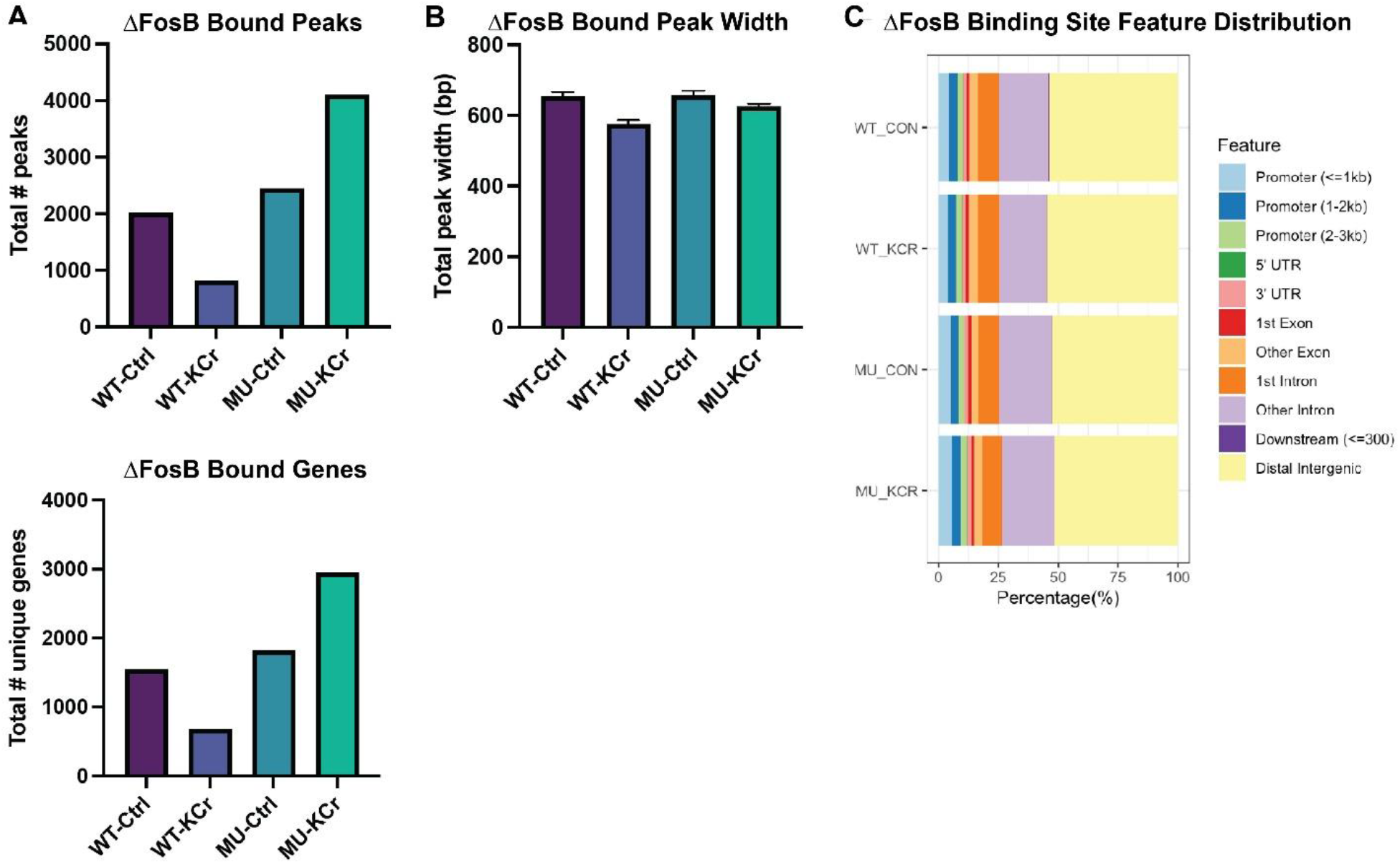
ΔFOSB C172 mediates oxidation-dependent loss of target gene binding in mouse hippocampus. **A)** CUT&RUN assay reveals the total number of ΔFOSB-bound peaks genome-wide in wild type (WT) and C172S (MU) mouse hippocampus after seven days of 25 mg/kg/day KCr in drinking water or normal control drinking water (Ctrl). **B)** Width of ΔFOSB-bound peaks. **C)** Distribution of ΔFOSB-bound peaks across genomic features. **D)** Total number of genes associated with ΔFOSB binding.

### ΔFOSB C172 oxidation mediates oxidative stress-induced cognitive dysfunction

We have previously shown that ΔFOSB/FOSB expression and function in hippocampus CA1 are necessary for cognition in mice, including spatial and reward learning [9, 11, 13, 14]. As it is well-established that oxidative stress is a key player in diseases of cognitive decline like AD [33, 34], we next sought to determine whether ΔFOSB/FOSB C172 oxidation plays a role in oxidative stress-mediated disruption of learning and memory. We first treated WT control mice with 25 mg/kg/day KCr in the drinking water for 7 days and examined baseline behaviors in the home cage. We found no differences in total movement or time spent feeding or drinking (**Fig S5**), indicating that this dose and timing regimen of KCr does not disrupt normal home cage behavior.

We next treated FOSB^C172S^ and WT littermate control mice with KCr and measured behaviors related to learning, stress, and reward. We found that KCr had no discernable effect on either genotype in elevated zero maze, cocaine CPP, or sucrose preference (**Fig 5A-C**; two-way ANOVA p>0.5 for genotype, KCr, and interaction effects), suggesting that this dose and timing regimen of KCr does not disrupt exploratory behavior or cause anxiety-like, mood, or reward dysfunction. To test learning and memory, we used the novel object location test, which we have previously shown is dependent on hippocampal ΔFOSB/FOSB function in mouse AD models [15]. We found that KCr treatment profoundly reduced the ability of WT mice to distinguish between objects left on their original location and objects that had been moved (**Fig 5D&E**), demonstrating oxidative stress-induced cognitive dysfunction. Strikingly, KCr treatment had no discernable effect on object location memory in FOSB^C172S^ mice (**Fig 5D&E**), demonstrating that ΔFOSB/FOSB C172 is a critical mediator of oxidative stress-induced cognitive dysfunction (5D: two-way ANOVA, genotype by drug interaction F=12.64, p<0.0001; 5E: one-way ANOVA, F=12.46, p<0.0001).

**Figure 5.**
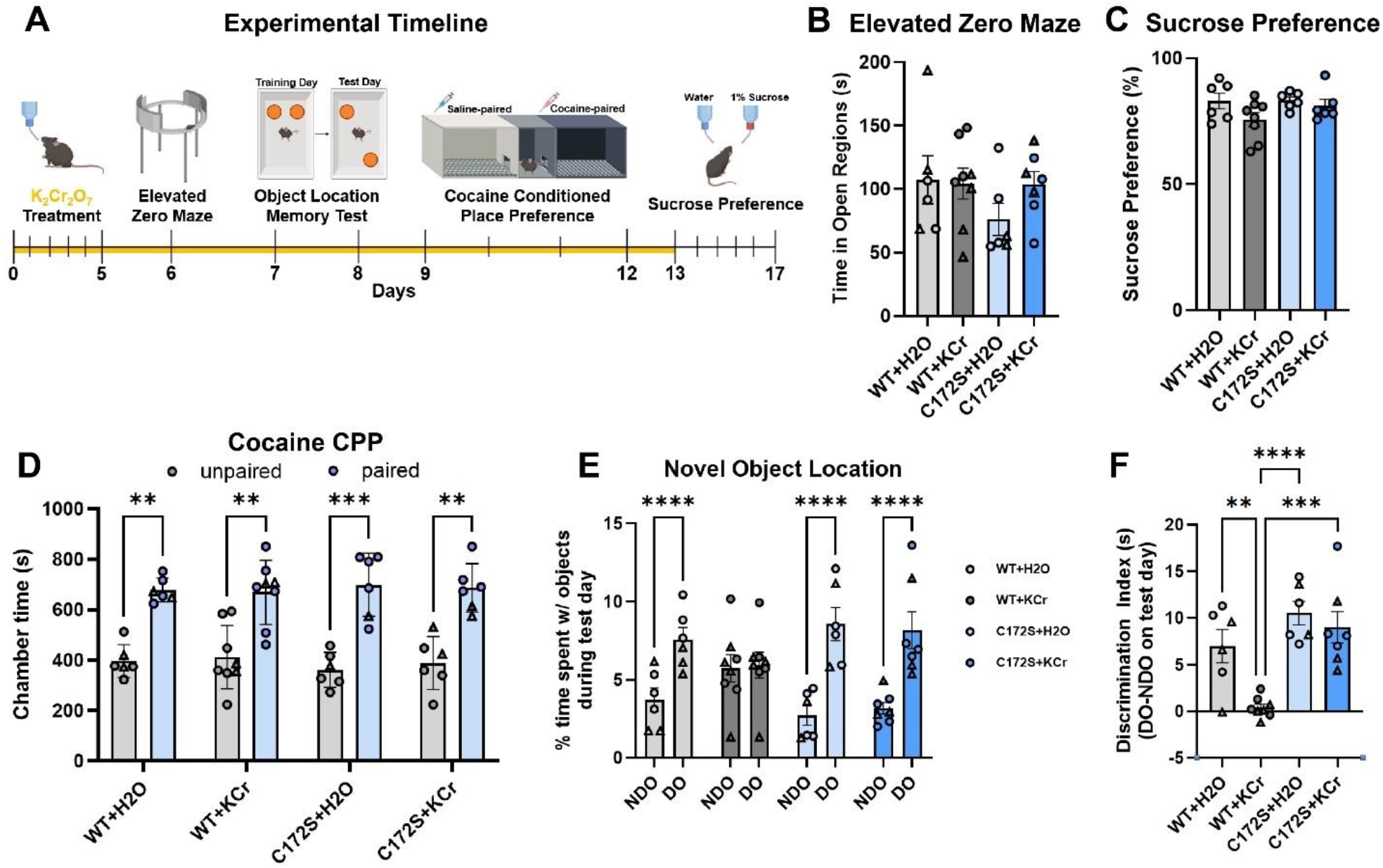
ΔFOSB C172 mediates oxidation-dependent cognitive dysfunction. **A)** Diagram of experimental timeline showing KCR treatment and order of behavioral tests (Biorender). Neither KCr treatment nor genotype had an effect elevated zero maze **(B)**, sucrose preference **(C)**, or cocaine CPP **(D). E)** KCr treatment prevented WT mice discriminating between displaced (DO) and non-displaced objects (NDO) in the novel object location test but had no effect on FOSB^C172S^ mice. **F)** Discrimination index in the novel object location test reveals that KCr treated WT mice had reduced object discrimination compared to all other groups. All graphs depict mean +/- SEM; **:p<0.01, ***:p<0.001, ****:p<0.0001.

## Discussion

This work demonstrates functional effects of oxidative stress driven by disulfide bond formation in an AP-1 transcription factor relevant to cognitive decline in a variety of diseases associated with redox imbalance. ΔFOSB expression and downstream regulation of target genes has been closely tied to AD in preclinical mechanistic studies, computational models, and postmortem analyses of patient brain tissue [10, 14, 15, 35]. Moreover, oxidative stress is a core pathological mechanism in AD [33, 34], and it is well established that altered neuronal function, and in particular synaptic dysfunction and loss, precede cell death and traditional histopathological markers of AD [36]. A key driver of this early AD neuronal pathophysiology is excitatory/inhibitory imbalance [37], particularly in the hippocampus, that is associated with seizure activity and mild cognitive impairment [38]. We have established that ΔFOSB functions to suppress hippocampal CA1 pyramidal neuron excitatory activity [11, 15, 39], and thus the inhibition of ΔFOSB function by oxidative stress-driven C172 disulfide bond formation we demonstrate here may act as a novel mechanism linking the redox imbalance in early stage AD to both seizure activity and cognitive decline.

Our recent work demonstrated that ΔFOSB can form disulfide bonds with its binding partner JUND *in vitro* using proteins purified from insect cells, and that the disulfide bond at C172 causes a “kink” in the structure of ΔFOSB that prevents its *in vitro* binding to AP-1 consensus sites in DNA [28, 30]. We were further able to demonstrate that oxidation-dependent redox-sensitive complexes containing ΔFOSB consistent with disulfide bonds to JUND could be formed in cells artificially overexpressing these proteins [30], but establishing that these complexes could form in the brain remained a challenge due to the lack of specific antibodies available for FOSB proteins. Thus, we created a transgenic mouse incorporating an HA epitope tag into the N-terminus of the *Fosb* gene, allowing us to purify ΔFOSB/FOSB-containing complexes from mouse brain under various redox conditions and probe them using existing ΔFOSB/FOSB antibodies without interference from non-specific bands. We were able to show that ΔFOSB/FOSB forms covalent, redox-sensitive complexes with molecular weights consistent with ΔFOSB-JUND dimers and ΔFOSB homodimers. We also observed higher molecular weight complexes consistent with ΔFOSB tetramers that we have observed *in vitro* [30]. Future experiments could use similar immunoprecipitates from these mice in combination with oxidative stress (i.e., KCr treatment), genetic AD models, or drug treatment, and mass spectrometry/proteomics, to determine the specific contents of these complexes and whether ΔFosB/FosB forms disulfide bridges with other proteins, besides JUND and itself, in the brain.

We further used CUT&RUN along with our novel ΔFOSB/FOSB C172S mutant mice to establish that ΔFOSB/FOSB oxidation at C172 inhibits binding to target genes in the mouse hippocampus *in vivo*. We found that treatment with KCr reduced ΔFOSB/FOSB binding to target genes in WT mice, but, surprisingly, we found that KCr significantly increased binding to target genes in the C172S mice. This could be explained by oxidation of other AP-1 family proteins preventing their binding to DNA, freeing up more AP-1 consensus sequences for the protected C172S ΔFOSB/FOSB complexes to bind. Indeed, FOS and JUN also display disulfide bond formation *in vitro* [26], and it is likely that oxidative stress in the brain prevents the binding of this complex to AP-1 sites, potentially liberating them for C172S ΔFOSB/FOSB-JUND binding.

We found that the specific dosing and timing of KCr that we used to induce protein oxidation in the brain caused impairment of object location memory in WT mice, and this effect was abrogated in the FOSB^C172S^ mice. Since this task is hippocampus-dependent, and because our previous work shows that ΔFOSB is critical for hippocampus-dependent learning [9-11, 13-15], the role of ΔFOSB oxidation in this form of learning fits our hypotheses well. We were surprised to find that the same dosing of KCr did not affect cocaine CPP or social interaction, both of which are dependent on ΔFOSB function in the nucleus accumbens and ventral hippocampus [9, 11, 12, 40]. It is possible that a higher dose of KCr or a longer period of exposure (beyond 7 days) would affect these behaviors in WT mice, and that the FOSB^C172S^ mice would be protected against such effects. However, higher doses and longer periods of oxidative stress cause strong neuronal dysfunction and death and have multiple deleterious effects on peripheral systems that reduce mobility, coordination, gastrointestinal function, and many other physiological issues that could cloud the results of behavioral tests measuring the rewarding effects of drugs or social interaction, so such experiments may require alternative approaches.

Nevertheless, it is very likely that ΔFOSB oxidation plays a role in physiology and behavior beyond cognition and spatial learning. Indeed, it is well established that cocaine and other psychostimulants induce oxidative stress in the brain and systemically [41, 42], suggesting that ΔFOSB oxidation could play a role in behavioral responses to drugs of abuse. Moreover, depressive disorders have been closely associated with oxidative stress [43], including mechanistic connections to the kynurenine pathway [44], and we and others have directly connected ΔFOSB expression and function in nucleus accumbens and hippocampus to susceptibility to anhedonia and social withdrawal in animal models [11, 12] and antidepressant action in mice and humans [12, 45, 46]. Thus, oxidative stress associated with mood disorders could drive some symptoms through changes in gene expression throughout the brain mediated by ΔFOSB C172 oxidation.

In order to take advantage of these new findings to treat diseases associated with oxidative stress, our group has been working to generate novel compounds that target ΔFOSB [7, 24, 28, 29]. We have published multiple series of compounds that regulate ΔFOSB binding to DNA *in vitro* and in cells, and we are currently engaged in screening these compounds in mouse models. Our previous work and the current data suggest that compounds directly targeting C172 through cysteine biochemistry could provide a potential mechanism to treat disorders directly associated with oxidative stress, but much work remains to ensure that such compounds do not interfere with the critical functions of ΔFOSB/FOSB in normal learning, motivation, or excitatory/inhibitory balance in the brain. Moreover, recent findings suggesting a role for ΔFOSB in non-neuronal cells in the brain [47, 48] and periphery [49, 50] make directly targeting ΔFOSB a challenge, but our work and that of many others justifies continued pursuit of ΔFOSB as an important therapeutic target.

In summary, the current work establishes the role of ΔFOSB C172 in regulating altered binding to gene targets driven by oxidative stress in both cultured cells and mouse brain and causally connects oxidation at this site to deficiencies in cognitive behavior caused by exogenous oxidative stress. Future studies may link these findings to pathology associated with AD and many other brain disorders that involve oxidative stress, potentially offering a new target for the treatment or prevention of these disorders.

## Notes

### Competing Interest Statement

The authors have declared no competing interest.

### Funding

- National Institute on Drug Abuse: DA040621
- National Institute of Mental Health: MH111604

## Citations

1. Uttara, B., et al., Oxidative stress and neurodegenerative diseases: a review of upstream and downstream antioxidant therapeutic options. Curr Neuropharmacol, 2009. 7(1): p. 65–74.

2. Cunha-Oliveira, T., A.C. Rego, and C.R. Oliveira, Oxidative Stress and Drugs of Abuse: An Update. Mini-Reviews in Organic Chemistry, 2013. 10(4): p. 321–334.

3. Cui, H., Y. Kong, and H. Zhang, Oxidative stress, mitochondrial dysfunction, and aging. J Signal Transduct, 2012. 2012: p. 646354.

4. Tonnies, E. and E. Trushina, Oxidative Stress, Synaptic Dysfunction, and Alzheimer’s Disease. J Alzheimers Dis, 2017. 57(4): p. 1105–1121.

5. Tan, M.G., et al., Genome wide profiling of altered gene expression in the neocortex of Alzheimer’s disease. J Neurosci Res, 2010. 88(6): p. 1157–69.

6. Robison, A.J. and E.J. Nestler, Transcriptional and epigenetic mechanisms of addiction. Nat Rev Neurosci, 2011. 12(11): p. 623–37.

7. Robison, A.J. and E.J. Nestler, DeltaFOSB: A Potentially Druggable Master Orchestrator of Activity-Dependent Gene Expression. ACS Chem Neurosci, 2022. 13(3): p. 296–307.

8. Hope, B.T., et al., Induction of a long-lasting AP-1 complex composed of altered Fos-like proteins in brain by chronic cocaine and other chronic treatments. Neuron, 1994. 13(5): p. 1235–44.

9. Gajewski, P.A., et al., Epigenetic Regulation of Hippocampal Fosb Expression Controls Behavioral Responses to Cocaine. J Neurosci, 2019. 39(42): p. 8305–8314.

10. Corbett, B.F., et al., DeltaFosB Regulates Gene Expression and Cognitive Dysfunction in a Mouse Model of Alzheimer’s Disease. Cell Rep, 2017. 20(2): p. 344–355.

11. Eagle, A.L., et al., Circuit-specific hippocampal DeltaFosB underlies resilience to stress-induced social avoidance. Nat Commun, 2020. 11(1): p. 4484.

12. Vialou, V., et al., DeltaFosB in brain reward circuits mediates resilience to stress and antidepressant responses. Nat Neurosci, 2010. 13(6): p. 745–52.

13. Eagle, A.L., et al., Experience-Dependent Induction of Hippocampal DeltaFosB Controls Learning. J Neurosci, 2015. 35(40): p. 13773–83.

14. You, J.C., et al., Epigenetic suppression of hippocampal calbindin-D28k by DeltaFosB drives seizure-related cognitive deficits. Nat Med, 2017. 23(11): p. 1377–1383.

15. Stephens, G.S., et al., Persistent ?FosB expression limits recurrent seizure activity and provides neuroprotection in the dentate gyrus of APP mice. Prog Neurobiol, 2024. 237: p. 102612.

16. Carle, T.L., et al., Proteasome-dependent and -independent mechanisms for FosB destabilization: identification of FosB degron domains and implications for DeltaFosB stability. Eur J Neurosci, 2007. 25(10): p. 3009–19.

17. Ulery-Reynolds, P.G., et al., Phosphorylation of DeltaFosB mediates its stability in vivo. Neuroscience, 2009. 158(2): p. 369–72.

18. Nestler, E.J., FosB: a transcriptional regulator of stress and antidepressant responses. Eur J Pharmacol, 2015. 753: p. 66–72.

19. Glover, J.N. and S.C. Harrison, Crystal structure of the heterodimeric bZIP transcription factor c-Fos-c-Jun bound to DNA. Nature, 1995. 373(6511): p. 257–61.

20. Yeh, S.Y., et al., Cell Type-Specific Whole-Genome Landscape of DeltaFOSB Binding in the Nucleus Accumbens After Chronic Cocaine Exposure. Biol Psychiatry, 2023. 94(5): p. 367–377.

21. Chen, J., et al., Chronic Fos-related antigens: stable variants of deltaFosB induced in brain by chronic treatments. J Neurosci, 1997. 17(13): p. 4933–41.

22. Hiroi, N., et al., Essential role of the fosB gene in molecular, cellular, and behavioral actions of chronic electroconvulsive seizures. J Neurosci, 1998. 18(17): p. 6952–62.

23. Jorissen, H.J., et al., Dimerization and DNA-binding properties of the transcription factor DeltaFosB. Biochemistry, 2007. 46(28): p. 8360–72.

24. Wang, Y., et al., Small molecule screening identifies regulators of the transcription factor DeltaFosB. ACS Chem Neurosci, 2012. 3(7): p. 546–56.

25. Nestler, E.J., Review. Transcriptional mechanisms of addiction: role of DeltaFosB. Philos Trans R Soc Lond B Biol Sci, 2008. 363(1507): p. 3245–55.

26. Abate, C., et al., Redox regulation of fos and jun DNA-binding activity in vitro. Science, 1990. 249(4973): p. 1157–61.

27. Yin, Z., Self-assembly of the bZIP transcription factor ΔFosB. Current Research in Structural Biology, 2020. 2: p. 1–13.

28. Yin, Z., et al., Activator Protein-1: redox switch controlling structure and DNA-binding. Nucleic Acids Res, 2017. 45(19): p. 11425–11436.

29. Kumar, A., et al., Chemically targeting the redox switch in AP1 transcription factor DeltaFOSB. Nucleic Acids Res, 2022. 50(16): p. 9548–9567.

30. Yin, Z., et al., Self-assembly of the bZIP transcription factor DeltaFosB. Curr Res Struct Biol, 2020. 2: p. 1–13.

31. Gajewski, P.A., G. Turecki, and A.J. Robison, Differential Expression of FosB Proteins and Potential Target Genes in Select Brain Regions of Addiction and Depression Patients. PLoS One, 2016. 11(8): p. e0160355.

32. Travacio, M., J. Maria Polo, and S. Llesuy, Chromium(VI) induces oxidative stress in the mouse brain. Toxicology, 2000. 150(1-3): p. 137–46.

33. Bai, R., et al., Oxidative stress: The core pathogenesis and mechanism of Alzheimer’s disease. Ageing Res Rev, 2022. 77: p. 101619.

34. Ionescu-Tucker, A. and C.W. Cotman, Emerging roles of oxidative stress in brain aging and Alzheimer’s disease. Neurobiol Aging, 2021. 107: p. 86–95.

35. Zachariou, M., E.M. Loizidou, and G.M. Spyrou, Topological influence of immediate-early genes in brain genetic networks and their link to Alzheimer’s disease. Comput Biol Med, 2025. 190: p. 110043.

36. Barthet, G. and C. Mulle, Presynaptic failure in Alzheimer’s disease. Prog Neurobiol, 2020. 194: p. 101801.

37. Vico Varela, E., G. Etter, and S. Williams, Excitatory-inhibitory imbalance in Alzheimer’s disease and therapeutic significance. Neurobiol Dis, 2019. 127: p. 605–615.

38. Vossel, K.A., et al., Epileptic activity in Alzheimer’s disease: causes and clinical relevance. Lancet Neurol, 2017. 16(4): p. 311–322.

39. Eagle, A.L., et al., DeltaFosB Decreases Excitability of Dorsal Hippocampal CA1 Neurons. eNeuro, 2018. 5(4).

40. Grueter, B.A., et al., ?FosB differentially modulates nucleus accumbens direct and indirect pathway function. Proc Natl Acad Sci U S A, 2013. 110(5): p. 1923–8.

41. Ng, F., et al., Oxidative stress in psychiatric disorders: evidence base and therapeutic implications. Int J Neuropsychopharmacol, 2008. 11(6): p. 851–76.

42. Sundar, V., et al., Psychostimulants influence oxidative stress and redox signatures: the role of DNA methylation. Redox Rep, 2022. 27(1): p. 53–59.

43. Bhatt, S., A.N. Nagappa, and C.R. Patil, Role of oxidative stress in depression. Drug Discov Today, 2020. 25(7): p. 1270–1276.

44. Sipahi, H., et al., The Interrelation between Oxidative Stress, Depression and Inflammation through the Kynurenine Pathway. Curr Top Med Chem, 2023. 23(6): p. 415–425.

45. Vialou, V., et al., Epigenetic mechanisms of depression and antidepressant action. Annu Rev Pharmacol Toxicol, 2013. 53: p. 59–87.

46. Robison, A.J., et al., Fluoxetine epigenetically alters the CaMKIIalpha promoter in nucleus accumbens to regulate DeltaFosB binding and antidepressant effects. Neuropsychopharmacology, 2014. 39(5): p. 1178–86.

47. Yutsudo, N., et al., fosB-null mice display impaired adult hippocampal neurogenesis and spontaneous epilepsy with depressive behavior. Neuropsychopharmacology, 2013. 38(5): p. 895–906.

48. Nomaru, H., et al., Fosb gene products contribute to excitotoxic microglial activation by regulating the expression of complement C5a receptors in microglia. Glia, 2014. 62(8): p. 1284–98.

49. Alim, M.A., et al., Glutamate triggers the expression of functional ionotropic and metabotropic glutamate receptors in mast cells. Cell Mol Immunol, 2021. 18(10): p. 2383–2392.

50. Duque-Wilckens, N., et al., FosB/DeltaFosB activation in mast cells regulates gene expression to modulate allergic inflammation in male mice. bioRxiv, 2025.

