## Supplemental Data and Methods for "Oxidation of ΔFOSB at Cys172 Controls Hippocampal Gene Targets and Learning"

### Supplemental Methods

*Recombinant protein expression and purification*

N-terminally hexaHis-tagged mouse ΔFOSB (MGHHHHHHAG followed by residues ($F^2$–$E^{237}$, a splice form of *Fosb*, accession number P13346) and mutants as well as N-terminally hexaHis-tagged mouse JUND (MGHHHHHH followed by residues ($E^2$-$Y^{341}$), accession number J04509) were expressed in Sf9 cells using baculovirus mediated overexpression (Bac-to-Bac system, Invitrogen) and purified using similar methods as we have described before [23, 24].

Wild-type $(His)_6$-ΔFOSB and the mutant $(His)_6$-ΔFOSB $C^{172}S$, were produced as homomers or as heteromers with $(His)_6$-JUND. Complexes of ΔFOSB/JUND or mutants were made either by combining the Ni-eluate of each protein in a 1:1 ratio, and then purifying the protein further by ion-exchange and gel filtration chromatography, or by co-infecting Sf9 cells with $(His)_6$-ΔFOSB (MOI ~1-1.5) and $(His)_6$-JUND baculoviruses (1:1 or 1:3 with respect to $(His)_6$-ΔFOSB) followed by further purification of the complex. Briefly, flash-frozen cell pellets from 3 L or 6 L insect culture were thawed, lyzed by sonication (in 20 mM Tris pH 8, 0.2% Triton X-100, 1 mM TCEP, 0.5 mM PMSF, 10 µg/ml leupeptin, 1 µg/ml pepstatin A), and then 300 mM NaCl, 5 mM $MgCl_2$ and 50 µg/ml DNAse added. In preparation for Ni-NTA chromatography (Invitrogen), the NaCl concentration was increased to 1M, 0.5 M NaBr and 5mM imidazole were added to the lysate, and the mixture subject to high speed centrifugation. Following Ni-NTA chromatography, the Ni-NTA eluate of $(His)_6$-ΔFOSB (wt or mutants) and $(His)_6$-JUND were mixed together (1:1 ratio) to form heteromeric complexes, diluted to 0.1-0.2 mg/ml and dialyzed overnight (in 25 mM Tris pH 9.0, 300 mM NaCl, 0.5% glycerol, 1 mM TCEP (or 5 mM DTT), 0.5 mM PMSF, 0.5% glycerol) and then dialyzed for 3 hr in a low salt buffer (25 mM Tris pH 9.0, 40 mM NaCl, 0.5% glycerol, 1 mM TCEP, 0.5 mM PMSF, 0.5% glycerol) before purification on a Resource-Q column (GE

Healthcare). For $(His)_6$-ΔFOSB (or mutant) homomeric complexes, ion-exchange was omitted, and instead the Ni-eluate was treated with 1 mM DTT, concentrated and then subject to size exclusion chromatography. Proteins were purified on a HiLoad 16/600 Superdex 75 pg column (GE Healthcare) equilibrated with 20 mM Tris pH 8, 1 M NaCl. For wild type and mutant ΔFOSB/JUND heteromers, additionally, fractions were confirmed to contain both proteins in a 1:1 complex based on SDS−PAGE prior to being pooled, concentrated and flash-frozen. $(His)_6$−ΔFOSB/$(His)_6$−JUND, $(His)_6$−ΔFOSB and variants with mutant $(His)_6$−ΔFOSB were stored in 20 mM Tris pH 8, 1 M NaCl at protein concentrations of 2 to 6 mg/ml. Protein purity was assessed by SDS-PAGE on 12% gels.

*Fluorescent polarization binding assay*

A fluorescence polarization (FP) based DNA-binding assay was performed as we have described before [23, 24]. Briefly, $(His)_6$-ΔFOSB, $(His)_6$-ΔFOSB $C^{172}S$, either as homomers or heteromers with $(His)_6$-JUND, were plated in a concentration series (0-800 nM) and a fixed concentration of TAMRA-labeled oligo TMR-*Cdk5* or TMR-*SCR* added (25 nM). The FP buffer for $(His)_6$-ΔFOSB/$(His)_6$-JUND heteromers was 20 mM HEPES pH 7.5, 180 mM NaCl and for $(His)_6$-ΔFOSB homomers, 20 mM HEPES pH 7.5, 65 mM NaCl with 1-5 mM TCEP (reducing conditions) or 10 μM diamide (oxidizing conditions) end concentration added. The protein:oligo samples were mixed and then manually dispensed in quadruplicates into 384-well round bottom low-volume black microtiter plates (Corning)  20 μl per well) and incubated for 15 minutes at room temperature. The FP signal was measured on a BioTek, Synergy Neo2 plate reader (excitation 530 nm, emission 590 nm) or using a Pherastar plate reader (BMG Labs; excitation 540 nm, emission 590 nm, 100 flashes per well); the target was set to 20 mP for each individual TMR-oligo by adjusting the gain on a well with oligonucleotide in the absence of protein). The FP signal observed for the free oligo (i.e., no protein) was set as the baseline value and subtracted from the FP values measured for oligos in the presence of varying amounts of protein.

The 19-mer *Cdk5* oligonucleotide ('*Cdk5* oligo') contains the forward and reverse strands of 5′-CGTCGGTGACTCAAAACAC-3′ (AP1 site underscored) from the AP1 site in the cyclin-dependent kinase 5 promoter. The non-specific DNA oligomer SCR (5'-GTATGCGATACGTCTTTCG-3') contains the same nucleotides as *Cdk5* but is scrambled. The oligos were made by annealing equivalent molar quantities of the complementary strands labelled with TAMRA at both 5′-ends (Sigma Aldrich) and heating them to 95 °C for 2.5 min, followed by slow cooling to room temperature (roughly 1 min/°C) and storage at -20°C as 50 μM stocks in annealing buffer (10 mM Tris pH 8, 50 mM NaCl).

*Animals*

All experiments were approved by the Institutional Animal Care and Use Committee at Michigan State University in accordance with AAALAC. Male C57BL/6J mice (3–5/cage, 7–8-week old and 25-30g upon arrival from Jackson Labs) were allowed at least 5d to acclimate to the facility prior to use and maintained at a constant temperature (22°C) and humidity (50-55%) on a 12 hour light/dark cycle. The hemagglutinin (HA)-tagged *Fosb* mouse line ($FOSB^{HA}$) was generated in house by the Nestler Lab at Mount Sinai and the FOSB C172S mutant mouse line ($FOSB^{C172S}$) was generated in the MSU Transgenic and Genome Editing Facility. Both lines were generated on a C57BL/6J background and backcrossed to C57/BL/6J wild type mice for five generations before used in experiments. All mice were group housed with *ad libitum* food and water at 22ºC and 50-55% humidity.

*Potassium Dichromate Treatment*

Mice were divided into four groups according to their genotype (WT or C172S) and treatment (water vs potassium dichromate). Two groups consisting of WT or C172S (control) received RO drinking water daily. The other two groups received a 25mg/kg dose of potassium dichromate ($K_2Cr_2O_7$) diluted in RO drinking water (0.11mg/mL). All four groups had access to either regular RO drinking water or $K_2Cr_2O_7$ drinking water *ad libitum*.

*Elevated Zero Maze*

The maze was constructed of grey acrylic in a circular track 5cm wide, 50cm in diameter, and elevated 50cm from the ground based on plans from ANY-maze (www.anymaze.com; Stoelting). The maze was divided into four quadrants of equal length with two opposing open quadrants with 1cm high acrylic curbs to prevent falls and two opposing closed quadrants with black acrylic walls 15cm in height. Animals were placed into the closed quadrant facing an open quadrant, and behavior was video recorded for 5 minutes. The percentage of time spent in the open quadrants was used as an assessment of anxiety-like behavior.

*Object Location Memory Test*

The object location memory test evaluates hippocampal-dependent memory and requires the mice remember the positions of two objects in a controlled cage arena. For the training trials, two identical objects were placed at adjacent corners of an arena that mice were allowed to explore over 3 trials of 3 minutes each, with 3-minute inter-trial intervals. In the testing trial, which occurred 24 hours after the last training trial, mice were placed back into the arena after one object had been displaced to another position. Mice that remember the original location of the two objects typically spend more time exploring the displaced object than the non-displaced object during the testing phase. Extra-arena spatial cues were used to help orient the mice during training and testing trials. The amount of time each mouse spent with each object in all trials was recorded by an observer blinded to genotype/treatment. The discrimination index was calculated as the difference between the time spent with the displaced object and the non-displaced object during the testing day.

*Cocaine Conditioned Place Preference*

CPP was conducted as previously described (Gajewski et al., 2019). Briefly, mice were tested for cocaine CPP in a 3-chamber CPP box (San Diego Instruments). On day 1, mice received a pre-test (no cocaine or saline) during which they were allowed to explore the entire box for 20 minutes. On days 2-3, mice received injections of saline paired with one chamber in the morning for 30 minutes and injections of cocaine (10.0mg/kg females; 12.5mg/kg males; IP) paired with the opposite chamber for 30 minutes in the afternoon (chamber counterbalanced by group). On day 4, mice were again tested in a post-test (no cocaine or saline) and allowed to freely explore the entire box.

*Two-bottle Choice for Sucrose Preference*

A standard two-bottle choice procedure was assessed across 5 days. Singly housed mice were first given access to two bottles of drinking water on the top of their home cage for 2 days to assess baseline drinking and to acclimate the mice to the bottles. Then, mice were given two-bottle choice, in which one of the bottles was replaced with a 1% sucrose solution for 3 days, each day alternating the position of the water and 1% sucrose solution. Sucrose solution consumed across days as a percentage of total liquid consumption was measured as an index of anhedonia.

*Immunoprecipitation*

Mouse brain was extracted from the hemagglutinin (HA)-mice and bilateral 12G dorsal hippocampal punches, centered on dentate gyrus, were collected in a fresh Eppendorf tube on ice. Tissue was harvested with 150 µL ice cold buffer (50mM Tric-HCl pH 7.5, 150M NaCl, 0.3% Tween) supplemented with protease inhibitor cocktail and phosphatase inhibitors (Sigma; St. Louis. MO); and treated with 1mM desired redox reagent (Dithiothreitol (DTT), N-ethylmaleimide (NEM), or Diamide) (Sigma; St. Louis, MO). Lysates were centrifuged at 4°C at 10,000 rcf for 10 minutes. 15 µL of supernatant were added to 5x laemmli buffer as the input Western blot sample. Remaining supernatant was separated from pellet and brought up to 800 µL with buffer in fresh Eppendorf tube. After 30 µL of 5/6,5mL Protein G Sepharose, Fast Flow beads (Sigma; St. Louis, MO) were washed 3x with 0.1% PBST, resuspended in 0.1% PBST and flipped in a vial rotator overnight with 3.5 µL anti-HA (Abcam ab 18181; Cambridge, MA) at 4°C in a fresh Eppendorf tube; the antibody-bound beads were washed with 0.1% PBST and the remaining supernatant from the mouse brain lysate was added. The Eppendorf tube was flipped overnight in a vial rotator at 4°C. 12 hours later, the Eppendorf tube containing the bead/lysate mix was spun down (1000rcf), and 50 µL of the supernatant was mixed with 20 µL 5x laemmli buffer for the flowthrough Western blot sample. Once the lysate/bead/antibody sample was washed 3x with 0.1% PBST, and the remaining supernatant was removed, immunocomplexes were eluted from the beads using 18 µL of 1.5x laemmli buffer (IP western blot sample). Before running, Western samples were boiled at 95°C for 1.5min. Procedure was repeated, with the exception that the Western blot samples were prepared using non-reducing laemmli buffer. This IP protocol resulted in the specific and effective precipitation of virtually all of the ΔFOSB in the lysate. The samples were run as indicated in the Western blot methods and probed for FOSB (1:500 FosB 5G4 2251s, antirabbit, Cell Signaling; Danvers, MA).

*Co-Immunoprecipitation*

Same protocol was used as in the immunoprecipitation methods; however, ∆FOSB binding partners were probed for during the Western blot (i.e. 1:500 JUND 5000s, antirabbit, Cell Signaling; Danvers, MA).

*Cell Culture and Transfection*

Neuro2a cells (N2a; American Type Culture Collection) were cultured in EMEM (ATCC; Manassas, VA) supplemented with 10% heat-inactivated fetal bovine serum (ATCC; Manassas, VA) and 5% Penstrep (Sigma; St. Louis) in a 5% $CO_2$ humidified atmosphere at 37°C. For the transient transfections with DNA, Neuro2A cell were seeded onto 12-well plates so as to reach 90-100% confluence the next day and were then transfected with the desired pcDNA3.1 plasmid using Lipofectamine 20000 (Promega; Madison, WI). A total of 1000ng of WT or mutant ∆FOSB or JUND was solely transfected, and 500ng of WT or mutant ∆FOSB and 1000ng JUND were co-transfected per well. ΔFOSB, FOS, and JUND cDNAs were obtained from our own pTetop-constructs [21], and subcloned into a pcDNA3.1 vector (Invitrogen; Carlsbad. CA). These pcDNA3.1-ΔFOSB/FOSB constructs were used for expression in mammalian cells and as a template for site-directed mutagenesis used to create mutant ΔFOSB. Recombinant HSV-ΔFOSB was prepared as described previously [29], and the preparation had a titer of $\sim 1 \times 10^8$pfu/ml.

*Site-directed mutagenesis*

Mutation of Cys172 and/or Cys222 to Ser within ΔFOSB in the pCDNA3.1 vector was accomplished using a Quick Change Site-Directed Mutagenesis kit (Agilent; Santa Clara, CA) and following the instructions of the manufacturer. The following mutagenesis primers were used:

**Cys172 to Ser:** 5' GACGGTTCCT***GGA***CTTAGCTGC 3' (forward primer)

5' GCAGCTAAG***TCC***AGGAACCGTC 3' (reverse primer)

**Cys222 to Ser:** 5' AGGGGATCTT***GCA***GCCCGGTTTG 3' (forward primer)

5' CAAACCGGGC***TGC***AAGATCCCCT 3' (reverse primer)

The mutated bases are in red, and the Cys codons are italicized and bolded.

*Intracellular protein extraction*

Approximately 48 hours post-transfection cells were washed with 1XPBS and detached from the plate using 0.3% Trypsin EDTA (Sigma; St. Louis, MO). Complete media was added to quench the trypsin and cells from each well were pipetted into a fresh Eppendorf tube. After cells were pelleted by centrifuging at 1000 rcf for 15minutes, the supernatant was removed and 100 µL RIPA buffer (10 mM Tris base, 150 mM sodium chloride, 1 mM EDTA, 0.1% sodium dodecyl sulfate, 1% Triton X-100, 1% sodium deoxycholate, pH 7.4) supplemented with protease inhibitors and phosphatase inhibitor cocktail with or without redox reagent (1mM DTT, NEM, or Diamide) was added. Lysate was sonicated at 30% amplitude 10x (1s on and 1s off). 50 µL of lysate was added to new Eppendorf tube with 15 µL reducing or non-reducing 5x laemmli buffer. Western blot samples were vortexed then boiled at 95°C for 1.5minutes. Western blot was performed (see methods).

*Cell Oxidation*

Neuro2a cells were plated into 12-well plates. 24 hours later (when cells were ~95% confluent) cells were transiently transfected with desired pcDNA3.1 plasmid containing WT or mutant ΔFOSB with or without JUND using Fugene 6. A total of 1000ng of WT or mutant ΔFOSB or JUND was solely transfected, and 500ng of WT or mutant ΔFOSB and 1000ng JUND were co-transfected per well. 1 hour post-transfection, cells were treated with Diamide or $H_2O_2$ containing media or ΔFOSB -targeting compound-containing media 24 hours post transfection oxidant-containing media was removed and new oxidant-containing media was added. Protein was extracted using the intracellular protein extraction and Western blots were run (see methods)

*Luciferase Assays*

Cells were plated into 12-well plates. Twenty-four hours later (when cells were ~95% confluent) cells were transiently co-transfected with a combination of 4×AP-1/RSV-Luc plasmid and pcDNA3.1 plasmid (Life Technologies; Carlsbad, CA) containing WT or mutant ΔFOSB with or without JUND Fugene 6. A total of 1000 ng of each DNA plasmid was transfected per well. Approximately 48 hours post transfection, cells were washed twice with 1 ml PBS and whole-cell lysates were prepared using 150 µl lysis buffer provided with ONE-Glo Luciferase Assay System (Promega; Madison, WI). 50 µl of the lysate were removed for Western blot analysis. The remaining lysates were incubated on ice for 5 minutes and the luciferase activity (luminescence) present in each sample was assayed using the substrates and protocol included in the ONE-Glo Luciferase Assay System. The luminescence of each sample was detected in triplicate using luminometer (TD-20/20 set at 2 s premeasurement delay and a 1 s measurement period). Luminescence was normalized to total ΔFOSB expression as assessed by Western blot.

*Western blotting*

Tissue punches and/or cell lysates were processed as above. After addition of Laemmli buffer, proteins were separated on 4-15% polyacrylamide gradient gels (Criterion System, BioRad; Hercules, CA), and transferred to PVDF membranes. Blots were probed using a polyclonal FOSB antibody (1:500 FOSB 5G4 2251s, anti-rabbit, Cell Signaling; Danvers, MA; referred to as "Pan-FOSB") and enhanced chemiluminescent detection (SuperSignal West Dura, Fisher Scientific; Waltham, MA). For **Fig 3A**, the putative ΔFOSB-specific antibody was also used (1:500 DeltaFOSB D3S8R, anti-rabbit, Cell Signaling; Danvers, MA). Films were scanned and band intensity was quantified using ImageJ software. The resulting histogram data were averaged for all samples of each group, and the lowest raw pixel intensity value was subtracted from all values for each graph, thus removing the background (which varied due to exposure time). For analysis of individual isoforms, a box was drawn around each of the pertinent bands (50kDa for FOSB; the 35-37kDa doublet for ΔFOSB) and average pixel intensity was calculated. A background value for each blot was generated from a region containing no specific bands and subtracted from all band values. Finally, membranes were stained for total protein with Swift Membrane Stain (VWR; St. Louis, MO) and all band values were normalized to total protein for each lane.

*Immunohistochemistry*

Adult male mice were terminally anesthetized (15% chloral hydrate) and transcardially perfused with PBS followed by 4% formalin. Brains were then postfixed overnight in formalin at 4°C and cryoprotected in 30% sucrose at 4°C until isotonic. Brains were sliced in 35 µm sections on a freezing microtome and immunohistochemistry for ΔFOSB expression was performed essentially as described previously [30]. Briefly, slices were blocked for 1hour in 0.3% Triton X-100 and 3% normal goat serum at room temperature then incubated overnight at 4°C in 1% normal goat serum, 0.3% Triton X-100, and primary antibody. Sections were washed, placed for 1 hour in secondary, and slices were mounted under glass coverslips for visualization on a fluorescent microscope. Images captured in both the red (FOSB) and green (GFP) channels. Fluorescent images were visualized on an Olympus FluoView 1000 filter-based laser scanning confocal microscope. Immunofluorescence was performed using the following primary antibodies: Anti-FOSB (1:500 FOSB 5G4 2251s, antirabbit, Cell Signaling; Danvers, MA), Anti-GFP (1:1000 GFP ab5450, antigoat, Abcam; Cambridge, MA). The following corresponding secondary antibodies were then used: Donkey anti-rabbit Cy3 (1:200, 711-165-152, Jackson Immunoresearch; West Grove, PA), Donkey anti-goat Alexa Fluor 488 (1:200, 705-545-147 Jackson Immunoresearch; West Grove, PA).

*Protein Carbonylation*

WT adult mice were treated with seven days of 25mg/kg/day KCr in drinking water or normal drinking water as a control, then sacrificed by cervical dislocation and decapitation. Hippocampus was harvested in 1 $mm^3$ punches and homogenized in 200 µl phosphate buffered saline with protease inhibitor cocktail and phosphatase inhibitors (Sigma; St. Louis. MO). Carbonylation was performed as per the manufacturer’s instructions (Protein CarbonylAssay Kit, Abcam, ab178020). Carbonylation was visualized by Western blot using the supplied anitbodies and chemiluminescent detection.

*CUT&RUN*

Nuclei preparation: nuclei were extracted from frozen hippocampal punches using a Dounce homogenizer to gently break down the tissue in 4 mL of lysis buffer. This buffer was composed of 320 mM sucrose, 5 mM $CaCl_2$, 0.1 mM EDTA, 10 mM Tris-HCl at pH 8.0, 1 mM DTT, 0.1% Triton X-100, 1.5 mM spermidine, and protease inhibitors. The homogenization involved 30 strokes with a loose pestle followed by 30 with a tight pestle. The homogenate was then filtered through a 40 µm strainer. A discontinuous sucrose gradient was prepared by layering 1.8 M sucrose, 10 mM Tris-HCl at pH 8.0, 1 mM DTT, 1.5 mM spermidine, and protease inhibitors under the lysate. Centrifugation was performed at 71,124 RCF for one hour at 4˚C, and the resulting nuclear pellet was resuspended in 1 mL of wash buffer containing 20 mM HEPES-NaOH (pH 7.5), 150 mM NaCl, 0.5 mM spermidine, 0.1% Triton X-100, 0.1% Tween-20, 0.1% BSA, and protease inhibitors, all maintained on ice. CUT&RUN: Nuclei were incubated with BioMag®Plus Concanavalin A (BP531, Bang Laboratories) for 10 min at room temperature. The beads were activated twice with 1.5 ml binding buffer (20 mM HEPES-KOH, pH 7.9, 10 mM KCl, 1 mM $CaCl_2$, and 1 mM $MnCl_2$). Next, the nuclei- bead complex were reclaimed at the magnet and incubated in antibody buffer containing rabbit anti-deltaFOSB (9890, Cell Signaling, 1:50 dilution) antibody at 4˚C overnight. The nuclei-beads complexes were then washed with antibody buffer and incubated with antibody buffer containing guinea pig anti-rabbit secondary antibody (ABIN101961, antibodies-online Inc., 1:100 dilution) at 4˚C for 1 h. Next, nuclei-bead complexes were washed with antibody buffer and incubated with antibody buffer containing pAGMNase (15-1116, EpiCypher) at 4˚C for 1 h. The beads were then washed with wash buffer twice and low-salt buffer (20 mM HEPES-NaOH, pH 7.5, 0.5 mM spermidine, 0.1% Triton X100, 0.1% Tween-20, and protease inhibitor) once before being incubated in calcium incubation buffer (3.5 mM HEPES-NaOH, pH 7.5, 10 mM $CaCl_2$, 0.1% Triton X-100, and 0.1% Tween-20) for 5 min on ice. The beads were reclaimed at the magnet and the reaction was stopped by addition of EGTA-stop buffer (170 mM NaCl, 20 mM EGTA, 0.1% Triton X-100, 0.1% Tween-20, 25 µg/ml RNase, and 20 µg/ml glycogen). DNA fragments were eluted by incubating the samples at 37˚C for 30 min with 500 rpm mixing. Next, the samples were centrifuged at 16,000 RCF for 5 min at 4˚C and the supernatant was reserved for DNA clean-up using NucleoSpin® Gel and PCR Clean-Up (740609, Takara) based on manufacture instructions. Library preparation: we used the NEBNext® Ultra™ II DNA Library Prep Kit for Illumina® (E7645, New England BioLabs) with NEBNext® Multiplex Oligos for Illumina® (Dual Index Primers Set 1) (E7600S, New England BioLabs) based on manufacture instructions. Libraries were submitted for sequencing by GENEWIZ on Illumina HiSeq 4000 machine with a 2 × 150-bp paired-end read configuration to a minimum depth of 25 million reads.

CUT&RUN Data Processing and Differential Binding Analysis: Sequencing data were processed using the NGSData-Charmer pipeline (GitHub commit DataCharmer). Adapter trimming was performed using Trim Galore (v0.6.5). Reads without detectable adapter sequences were secondarily trimmed with Cutadapt (v2.10) to remove 6 base pairs from the 3' end. Trimmed reads were aligned to the mouse reference genome (mm10) using Bowtie2 (v2.4.1) with the parameters -dovetail --phred33. Duplicate reads were removed using Samtools (v1.10) rmdup. For visualization, TDF files were generated using IGVtools (v2.5.3), and bigwig files were created using Deeptools (v3.5.0) (bamCoverage --binSize 10 --normalizeUsing RPKM). Merged bigwig files were generated from BAM files using Samtools merge and index, followed by normalization with Deeptools (bamCoverage --extendReads -ignoreDuplicates --centerReads --normalizeUsing CPM --binSize 10). Peaks were called for each replicate and its IgG control using MACS2 (v2.2.6)120 (macs2 callpeak -f BAMPE --pvalue 0.05 --keep-dup) and filtered using IDR (v2.0.4.2)121 at a 5% threshold. High-confidence peaks were merged using Bedtools (v2.29.2)122. Peak annotation was performed using ChIPseeker (v1.22.1)123 in R (v3.6.1). Peak overlap analysis was conducted with HOMER (v4.9) using the mergePeaks module. To compare CUT&RUN binding profiles, differentially bound regions were identified using the DiffBind package (v2.10.0) in R (v3.5.3), applying thresholds of $|\log_2$ fold change$| \geq 1$ and FDR $< 0.05$. Consensus peaksets were derived from reproducible peaks across conditions. Peak regions were annotated to their nearest gene using HOMER67 annotatePeaks.pl, and enriched motifs were identified with findMotifGenome.pl using scrambled sequences as background controls.

*Generation of Fosb$^{HA}$ mice*

The HA-*Fosb* mouse strain was generated by CRISPR/Cas9-mediated genome editing in C57BL/6N embryonic stem (ES) cells. A donor template containing a 27-bp hemagglutinin (HA) epitope tag sequence (encoding the amino acids YPYDVPDYA) was designed for insertion into the *endogenous Fosb* locus between the translational start codon (ATG) and the codon for the second amino acid (phenylalanine, TTT), in frame with the coding sequence. Cas9 nuclease and a guide RNA targeting the 5′ region of the *Fosb* gene were co-electroporated with the donor template into ES cells. Correctly targeted clones were identified by PCR screening of the 5′ and 3′ homology junctions and confirmed by Sanger sequencing to ensure proper in-frame insertion of the HA tag and absence of indels at the non-edited allele. Targeted ES cell clones with a normal karyotype were injected into blastocysts to generate chimeric mice. Germline transmission was confirmed by genotyping, and the line was backcrossed to C57BL/6J for at least five generations prior to experimental use. Genotyping was performed by tail DNA PCR using the following primers:

95624cof-NES1: 5' - CTG AGC TAA GTG GGA AGG AGA GGC TTC - 3'

95625cof-NES1: 5' - CGA GAG GGG TAT AAT CAG TCT AGG TGG - 3'

Example PCR gel electrophoresis bands can be found in **Fig S1B**.

*Statistical Analysis*

All analysis was performed using Prism software (GraphPad). Student's *t* tests were used for all pairwise comparisons (indicated in Results where the *t* value is given), and one-way or two-way ANOVAs were used for all multiple comparisons (indicated in Results where the *F* value is given), followed by Bonferroni or Tukey *post hoc* tests where appropriate.

**Supplemental Figure Legends**

**Figure S1: Generation of FOSB$^{HA}$ Mouse. A)** Schematic of strategy for generation of a transgenic mouse with an insertion encoding the HA tag on the N-terminus of the *Fosb* open reading frame. **B)** Example gel showing PCR product of genotyping reaction from a wild type (WT) mouse and both heterozygous and homozygous FOSB$^{HA}$ mice.

**Figure S2: Generation of FOSB$^{C172S}$ Mouse. A)** Schematic showing gRNA targeting the DNA encoding cysteine 172 in exon 3 of the *Fosb* gene. This gRNA introduced a single base pair mutation of the TGC codon encoding C172 into an AGC codon encoding serine. **B)** Sanger sequencing plot demonstrating the introduction of the T to A mutation in the first germline mouse. All subsequent genotyping was performed by similar Sanger sequencing.

**Figure S3: FOSB$^{C172S}$ Mice have normal baseline behavior and characteristics. A)** FOSB$^{C172S}$ mice do not differ from wild type littermates (WT) in time spent in the open or closed arms of the elevated plus maze. WT and FOSB$^{C172S}$ mice do not differ in sucrose preference **(B)**, ability to discriminate the displaced object in the novel object location test **(C)**, or in preference for the cocaine-paired chamber in a conditioned place preference test **(D)**. All graphs depict mean +/- SEM; **:p<0.01, ***:p<0.001.

**Figure S4: KCr treatment increases oxidation in mouse brain. A)** Western blot demonstrating that treatment of wild type mice with seven days of 25mg/kg/day KCr in drinking water had no effect on ΔFOSB protein expression in the hippocampus compared to mice with normal drinking water. **B)** Protein carbonylation Western blot demonstrates increased carbonyl groups introduced into proteins by oxidative reactions in the hippocampus of mice with seven days of 25mg/kg/day KCr in drinking water compared to mice with normal drinking water.

Figure S1

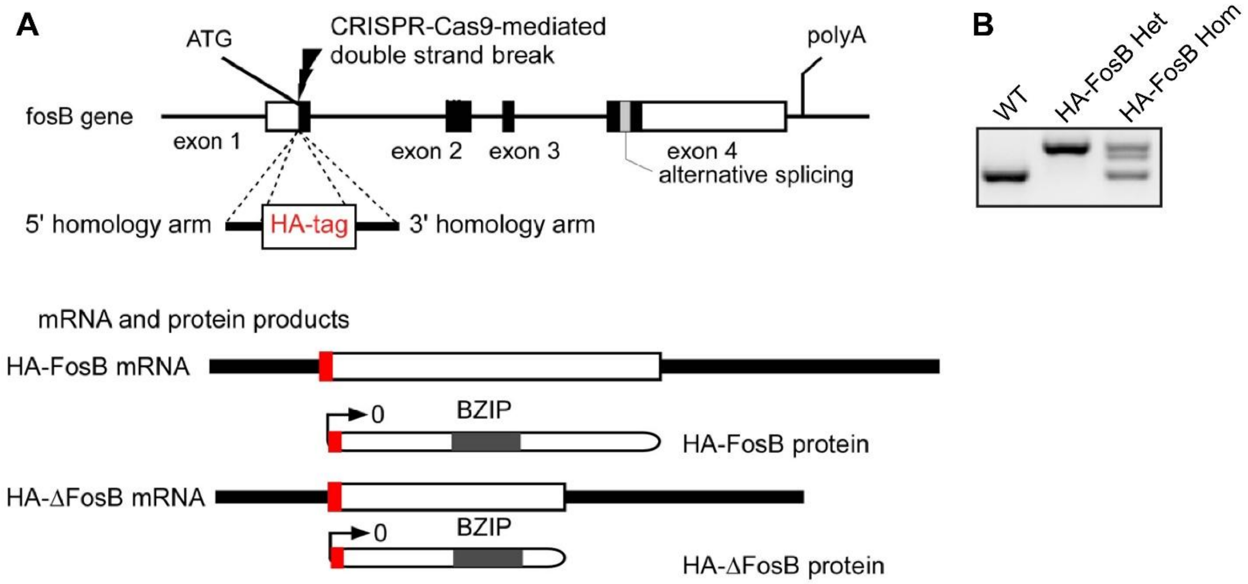

Figure S2

**A** Mutation of TGC encoding cysteine 172 to AGC encoding a serine

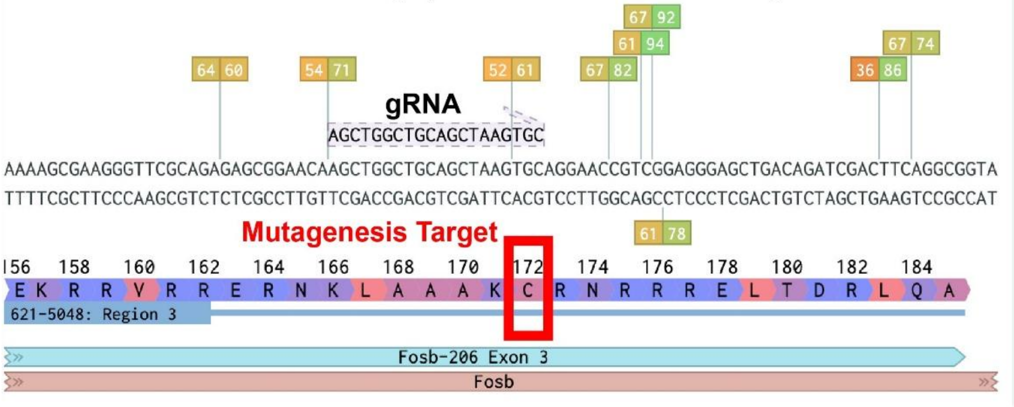


**B** Sequencing demonstrating selective T→A mutation in germline

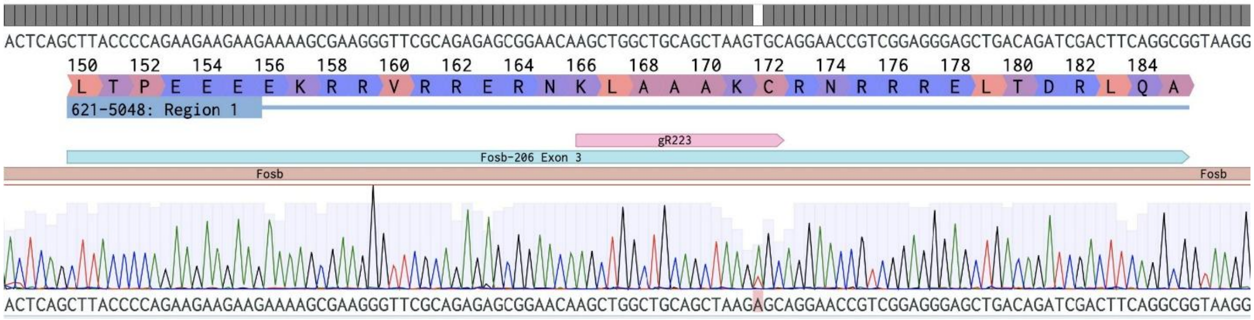

Figure S3

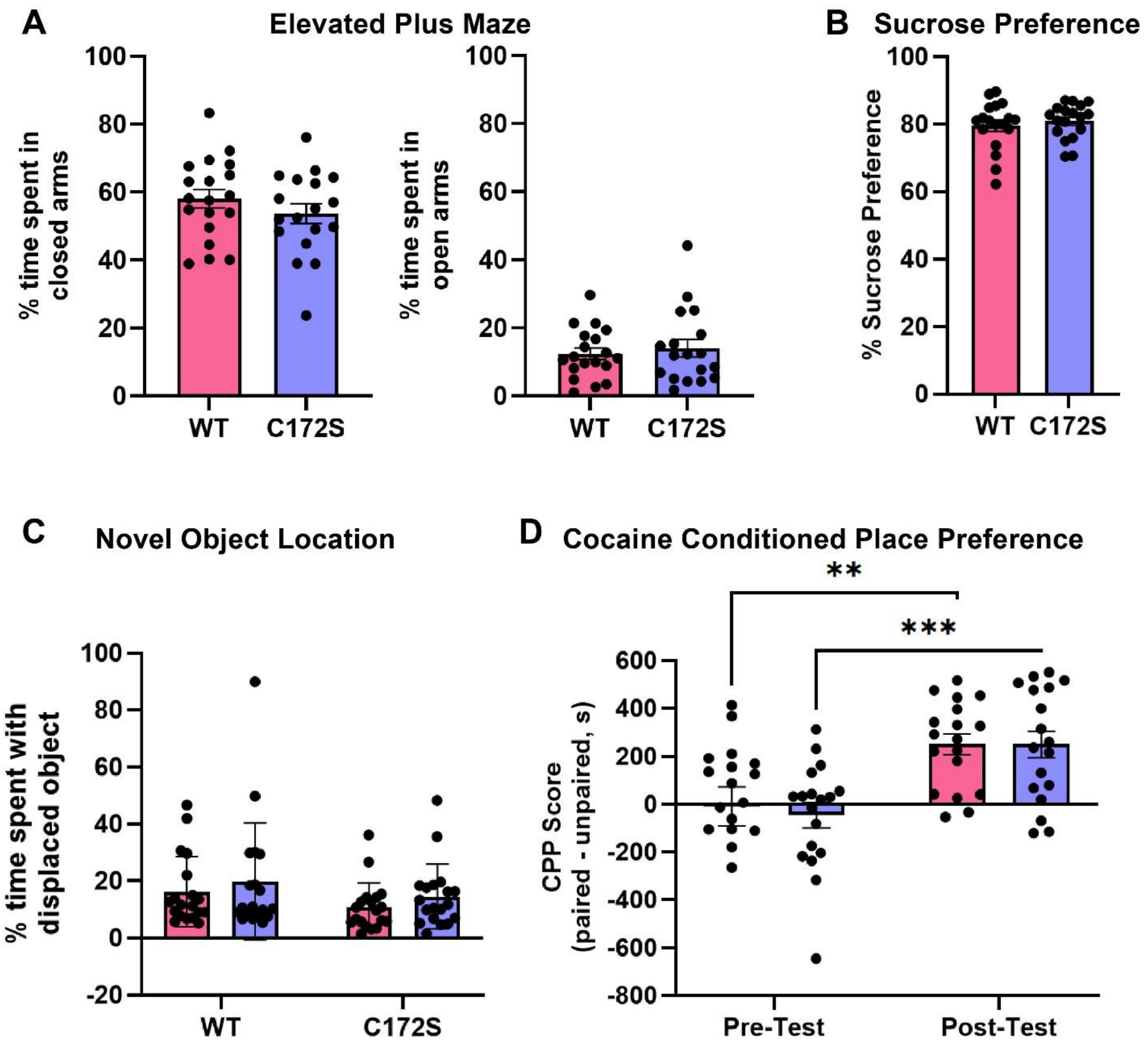

A
Elevated Plus Maze
% time spent in closed arms
% time spent in open arms
100
80
60
40
20
0
WT
C172S
B
Sucrose Preference
% Sucrose Preference
C
Novel Object Location
% time spent with displaced object
-20
D
Cocaine Conditioned Place Preference
**
***
CPP Score (paired - unpaired, s)
600
400
200
-200
-400
-600
-800
Pre-Test
Post-Test

Figure S4

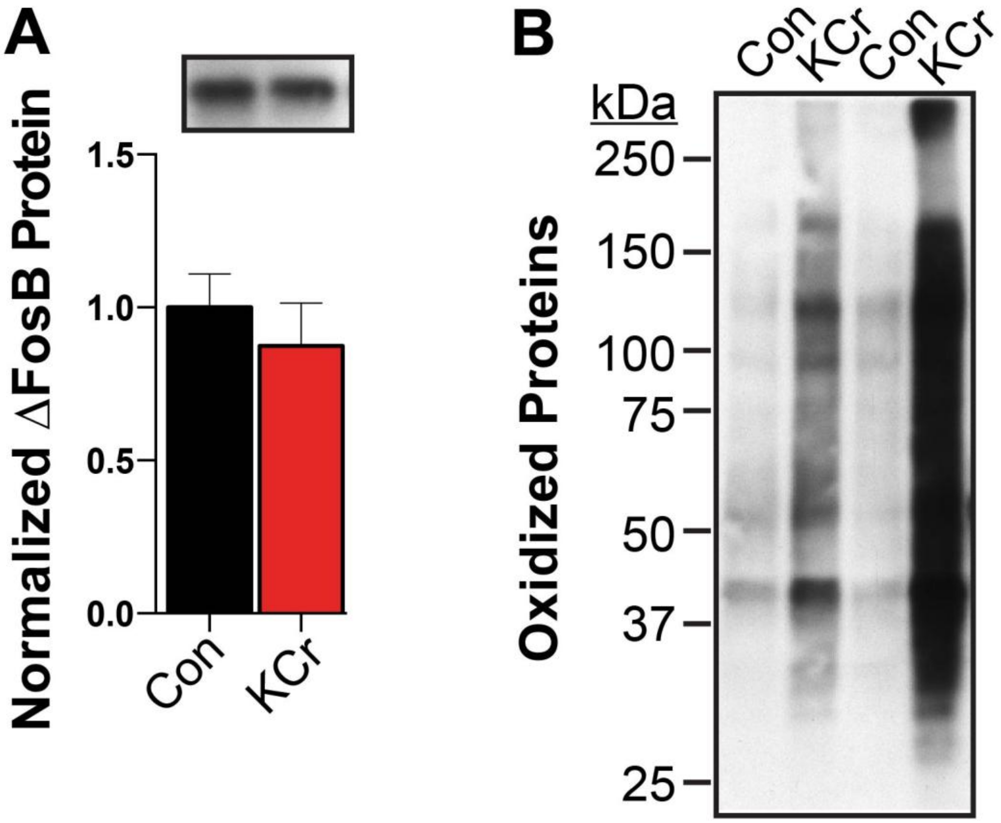

A
Normalized ΔFosB Protein
1.5
1.0
0.5
0.0
Con
KCr
B
Con
KCr
Con
KCr
kDa
250
150
100
75
50
37
25
Oxidized Proteins
